# Pseudozyma aphidis bio-active extract inhibit plant pathogens and activate induce resistance in tomato plants

**DOI:** 10.1101/2024.12.18.629071

**Authors:** Raviv Harris, Maggie Levy

## Abstract

The constant growth in the world population demands a constant increase in agricultural yields. One of the main ways to increase agricultural yields is by improving the control of pests and pathogens. Human health and environmental concerns regarding the traditional synthetic pesticides challenge the scientific community to discover new and less harmful ways to control pests, such as the development of biocontrol agents and natural-based pesticides.

Previous studies have established that application of live *P. aphidis* can be used for biocontrol of diverse fungal and bacterial phytopathogens. Here demonstrate activity of two semi-purified fractions from *P. aphidis*, one containing the antimicrobial metabolites and the other containing the resistance inducing metabolites. Our results from the *in vitro* experiments with the antimicrobial extract show that *P. aphidis* metabolites strongly inhibit important fungal and bacterial phytopathogens. *In planta* experiments demonstrated a significant, dose-dependent reduction in disease infection when a spore suspension of *B. cinerea* was treated or exposed to the extracted metabolites. From the other hand, our results showed that the application a semi-purified aqueous fraction from *P. aphidis* on tomato plants rapidly up-regulated the expression of defense-related genes, which are associated with both the induced systemic resistance and the systemic acquired resistance pathways.

In conclusion, this study further enhances our understanding of the biochemical mechanisms behind *P. aphidis* main modes of action: antibiosis and induced resistance. It also demonstrates the great potential of this unique biocontrol agent as a source for new natural-based pesticides and/or enhanced resistance substances.

## Introduction

Pathogenic microorganisms and pests attack agricultural crops and induce diseases that reduce the plant’s growth rate and yield. Constant use of chemical-based pesticides allows farmers to control the spread of pathogens and pests and help ensure stable and prosperous agricultural systems (Denholm & Rowland, 1992a, Leroux *et al*., 2002). However, concerns over the potential impact of pesticides on the environment have now become more pressing. Recent, more stringent pesticide registration procedures and regulations have reduced the number of synthetic pesticides available in agriculture. A possible replacement to the traditional synthetic pesticides is the discovery and developing of natural product-based pesticides, which are less harmful to human health and for the environment (Dayan *et al*., 1999, Dayan *et al*., 2009, Copping & Duke, 2007, Kim *et al*., 2017, Dhankhar & Kumar, 2023, Donley *et al*., 2024, Daraban *et al*., 2023, Zhang *et al*., 2023, Singh & Maurya, 2023).

Along with plants, microorganisms are a potential source of such biological pesticides. These compounds may either be used directly as natural pesticides, or as lead molecules for the develop of commercially synthetic pesticides (Balba, 2007, Swapan *et al*., 2024, Zhang et al., 2023). Since antibiosis is a well-established mechanism of biocontrol, microorganisms which are already known as biocontrol agents (BCAs) are more likely to produce natural pesticides (Raaijmakers *et al*., 2002, Tran *et al*., 2023).

*Pseudozyma aphidis* (isolate L12) found to have biological activity against diverse plant pathogens using multiple mode of actions includes secretion of antibiotic compounds that act as pesticides and parasitism (Buxdorf *et al*., 2013a, Buxdorf *et al*., 2013b, Barda *et al*., 2015). Additionally to the direct control of phytopathogens (via parasitism and antibiosis), *P. aphidis* isolated in our laboratory was found to indirectly protect plants by local and systemic induction of SAR and ISR, and priming the plant defense machinery to induce stronger defense activation after pathogen infection (Buxdorf et al., 2013a, Barda et al., 2015).

Some beneficial microbes, especially biocontrol agents and growth–promoting bacteria and fungi in the rhizosphere, can as part of other mode of actions, induce resistance in plants. Beneficial microbes are generally associated with induced systemic resistance (ISR) pathway that involve the phytohormones Jasmonic acid (JA) and ethylene (ET) signaling pathway. Nevertheless, they can also activate the systemic acquired resistance (SAR) which associate mainly with the phytohormones salicylic acid (SA) signaling pathway, but can also be independent of both pathways (Buxdorf et al., 2013, Pieterse et al., 2014; Salwan et al., 2023; Garcia-Montelongo et al., 2023). Several fungal BCAs, mainly of the genus Trichoderma, have been reported to elicit ISR and priming in colonized plants (Rodrigues et al., 2023; Salwan et al., 2023; Perazzolli et al., 2011; Yoshioka et al., 2012; Molitor et al., 2011; Aime et al., 2013).

Prolonged activation of inducible defenses involves major costs that affect plant growth and reproduction. Therefore, for the ISR to be affordable and beneficial, it should only be activated when the plant is exposed to attack (Heil, 2002, Agrawal, 1998, Baldwin, 1998). To overcome this, beneficial microbes induce a unique physiological state called priming, which causes the plant to respond with faster and stronger activation of the induced defense responses, but only following an attack by pathogens (Conrath et al., 2006; Honig M et al., 2023; Harris CJ et al., 2023).

In this work we demonstrate that bio active metabolites extracted from the biocontrol agent *P. aphidis* (isolate L12) can also inhibit a wide range of fungal and bacterial phytopathogens, both in vitro and in planta and can activate induced resistance in tomato plants.

## Materials and methods

### P. aphidis culture

*Pseudozyma aphidis* (isolate L12 collected in Israel) maintained on solid culture, PDA (Difco), at 25°C and transferred to fresh medium monthly. Liquid cultures were maintained in PDB (Difco) for 7 to 10 d at 27°C, on a rotary shaker, at 150 RPM. We obtained 10^8^ cells mL^−1^ after 5 to 7 d in liquid culture.

### Propagation of plants and pathogens

*Botrytis cinerea* (B05.10)( 22°), *Fusarium oxysporum f. sp. Lycopersici* (27°) and *Alternaria alternata* (25°) were grown on PDA medium, under 12-h daily illumination. *Rhizoctonia solani*(27°) and *Pythium*(27°) were grown on PDA medium, in dark.

*Clavibacter michiganensis subsp.michiganensis* strain CMM44 (30°C), *Xanthomonas campestris pv. vesicatoria* (28°C), *Agrobacterium tumefaciens* (32°C), *Erwinia amylovora* (37°C), and *Pseudomonas syringae pv. tomato* (27°C) (from our private collection) were grown on nutrient agar medium (NA; Difco) in a controlled incubator, in complete darkness, at the indicated temperatures.

We used wild-type *Solanum lycopersicum* (cv. Micro-Tom) plants. They kept in a greenhouse, at a constant 26°C, under natural light conditions.

### Bio-active compounds extraction from *P. aphidis* biomass

Erlenmeyer flasks, each with a working volume of 1.5 liter of PDB medium, were inoculated with 3-cm^2^ blocks of PDA carrying mycelia and/or spores of *P. aphidis* and incubated for 7 days, at 27°C, in dark, with constant agitation at 150 rpm. The fungal cells were spun down (Sorvall RC 5C plus) and extracted with several solvents (Distilled water (DW), Methanol (meOH), Acetone, Ethyl acetate (EtAc), Chloroform and Hexane), at 60°C. The extracted fraction was collected, filtered and concentrated using a rotor evaporator (Buchhi, Flawil, Switzerland) at 42°C. All extracts were subjected to bio-activity assays against *B. cinerea* and *A. tumefaciens*, using the agar diffusion method (described in further details below).

### Isolation of a semi-purified aqueous fraction from *Pseudozyma aphidis* isolate L12 extract, containing the resistance inducing metabolites as main components

*P. aphidis* extract was mixed with an equal portion of water and the mixture was stirred in a Pyrex cylinder for 5 minutes. After phase separation, the aqueous fraction was collected. New water was added to the mixture and the process was repeated two more times. The collected aqueous fractions were combined, completely dried and re-dissolved in water for complete elimination of any traces of organic solvents from the fraction.

### In vitro inhibition of plant pathogens

Germination inhibition of fungi and growth inhibition of bacteria were evaluated using the agar diffusion method. 20µl of a diluted bio-active extract (EtOAc), corresponding to 1mg in dry weight was aliquot on Whatman paper discs (6 mm diameter). The discs were placed in the center of PDA plates, embedded with plant pathogens .For fungi inhibition, a spore suspension of 2×10^5^ spores mL^−1^ was mixed in 50ml of PDA and equally distributed between 5 petri dishes. For bacteria inhibition, 1ml of bacteria suspension with optical density of 1.0 O.D, at 600nm was mixed in 50ml of PDA and equally distributed between 5 petri dishes (9cm diameter). The area of the inhibition zone was measured after 24 h incubation at the same temperature (depicted in parenthesis in the “Propagation of plants and pathogens” section) that was used for the pathogen’s propagation.

In order to evaluate the antifungal activity of *P. aphids* extract on the colony mycelial growth, fungal mycelial plugs (4mm diameter) were obtained from mature colony edges and placed at the center of 9cm Petri dishes, containing PDA culture media, supplemented with increasing concentrations of the extract. The cultures were incubated at 25 ± 2°C. When the colony of the control group reached close to the confluence of the Petri dish, all the petri dishes were photographed and the area of the colonies was measured using ASSESS 2.0, image analysis software for plant disease quantification (APS Press, St. Paul, MN, USA). The relative growth inhibition of each treatment compared to control was calculated as % of inhibition, using the following formula:

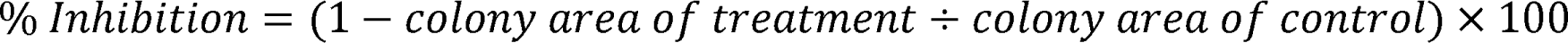

The IC50 value of the extract against each fungus was calculated from the trend line of the correspondence scatter plot (% inhibition to concentration), using the formula:

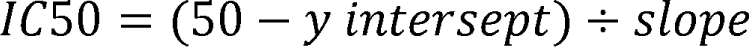

### Inhibition of *B. cinerea* on tomato plants

Tomato leaflets were detached and placed in a closed tray, on water-agar media to conserve the freshness of the leaflets. each leaflet was inoculated with a 4µl drop of inoculums, composed of 50% grape juice, 45% aqueous spore suspension (1500 spores/µl) and 5% of *P. aphidis* extract (0.35mg/ml, 0.7mg/ml, 1.4mg/ml, 4.2mg/ml, 7mg/ml and EtOAc for control). After 96 hours, the leaflets were photographed and the area of the lesions was measured using ASSESS 2.0, image analysis software.

For evaluation of *B. cinerea* growth inhibition on whole tomato plants, we inoculated 15-30 leaflets of each plant in the same method and inoculums as the detached leaflets. The plants were kept in high humidity chambers, at 24°. After 120 hours, the inoculated leaflets were detached and photographed, and the area of the lesions was measured using ASSESS 2.0, image analysis software.bi

In order to evaluate the preventive effect of the extract, 15-30 leaflets of each plant were sprayed with 1% TWEEN80 solution, mixed with increasing concentrations of the extract (0.2mg/ml, 1mg/ml, 5mg/ml, 10mg/ml and 15mg/ml). Two hours later, the sprayed leaflets were inoculated with *B. cinerea* (2000 spores/leaflet). The disease symptoms were evaluated after 7 days as the percentage of infected leaflets/plant, using the formula:

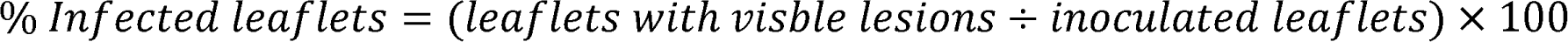

### Thin Layer Chromatography of antimicrobial compounds from *P. aphidis* extract

Chemical analysis of *P. aphidis* extract was conducted using Thin Layer Chromatography (TLC). TLC were carried out by spotting 10 mg extract, corresponding to 50 µl of the extract on TLC plates (10 x 20 cm, Silicagel 60 F_254_, Merck, Germany), 2 cm from the bottom. TLC plates were developed under saturated conditions with the following solvent system: Chloroform/Methanol/Water (75/25/2.5). After migration, TLC plates were air dried and observed under UV light (UV-A: 354 nm ; UV-B: 302 nm and UV-C: 254 nm) in order to visualize the presence of metabolites on plates. The metabolites are characterized by their Rf (Retention factor) corresponding to: Rf = migration distance of the compound/migration distance of the solvent. After observation under UV lights, TLC plates were covered with a calibrated conidia suspension of *B. cinerea,* prepared in V8-Agar medium (pH 4.6). The TLC plates were then incubated at 25°C, in darkness, for 7 days. Metabolites with antifungal activity exhibit inhibition spots on TLC plates.

### Induced resistance activation

For local defense induction, tomato plants were completely and evenly sprayed with the semi-purified aqueous fraction in concentration of 20 mg ml^-1^ (DW was used as control). The plants were sampled after: 12, 24, 72 and 120 h. Sampling of the plants was performed by cutting a small piece of the edge of three leaflets, each from a separate plant (in total, about 50 mg of tissue).

For systemic defense induction, the outmost leaflet of three leaves were treated with six droplets of 5 µl of the semi-purified aqueous fraction in concentration of 100 mg ml^-1^ (DW was used as control). The plants were sampled after: 12, 24, 72 and 120 h. Sampling of the plants was performed by cutting a small piece of the edge of three leaflets from untreated distal tissue of a separate plant (in total, about 50 mg of tissue). Sampling in each time point was performed in triplicates (designated as dependent biological repeats). In addition, the entire process was repeated with sampling of leaflets from un-treated leafs, to obtain differentiation of two levels of systemic effect: short distance systemic effect and long-distance systemic effect.

### Priming of defense-related genes

For local priming, tomato plants were completely and evenly sprayed with the semi-purified aqueous fraction in concentration of 20 mg ml^-1^ (DW was used as control). 24 h post treatment, 5 leaflets of each plant were inoculated with a 4 µl drop of *B*. *cinerea* inoculums, composed of 50% grape juice and 50% aqueous spore suspension (50 spores µl^-1^). The plants were sampled 24 and 48 h post the inoculation.

For systemic priming, the outmost leaflet of three leafs was treated with six 5 µl drops of the semi-purified aqueous fraction in concentration of 100 mg ml^-1^ (DW was used as control). 48 h post treatment, two other leaflets of each of the treated leaves were inoculated with a 4 µl drop of *B. cinerea* inoculums, composed of 50% grape juice and 50% aqueous spore suspension (50 spores µl^-1^). The plants were sampled 24 and 48 h post the inoculation.

### RNA isolation

RNA was extracted from the 50 mg lysed plant tissue with TRI-Reagent (Sigma-Aldrich) and purified with Total RNA Mini Kit (Geneaid), followed by treatment with TURBO DNA-free (Invitrogen) to remove genomic DNA contamination.

### Quantitative reverse transcription PCR analysis

Total RNA (1 μg) was reverse transcribed with High Capacity cDNA Reverse Transcription Kit (Applied Biosystems). Quantitative reverse transcription PCR was performed with the SYBR master mix and StepOne real-time PCR machine (Applied Biosystems). The thermal cycling program was as follows: 95°C for 20s and 40 cycles of 95°C for 3s and 60°C for 30s. Relative fold change in gene expression, normalized to *GAPDH* and *TIP41*, was calculated by the comparative cycle threshold 2^-ΔΔCt^ method. Primers used in the qRT-PCR analysis are listed in **Table S1**.

### Statistical analysis

Statistical analysis was conducted with GraphPad Prism software. For multiple comparisons, either one-way ANOVA and Dunnett’s multiple comparisons test, or two-way ANOVA (Bonferroni’s multiple comparisons test) were used according to the experiment. Significance was accepted at P < 0.05.

## Results

### Extraction and isolation of two semi-purified fractions from P. aphidis isolate L12

Our hypothesis was that *P. aphidis* secrets different sets of metabolites to accomplish two of its main modes of action: antibiosis and induced resistance. To examine this hypothesis, *P. aphidis* bioactive metabolites had to be efficiently produced, extracted and divided into two distinct fractions according to their biological activity.

### Bioassay-guided isolation of a semi-purified fraction, containing the antimicrobial metabolites as main components

Biologically active metabolites were extracted from *P. aphidis* with the following organic solvents: EtOAc, methanol, heptane, hexane, MTBE, DCM, Acetone and acetonitrile. Extracts were determined for their antimicrobial activity against *B. cinerea* and *A. tumefaciens* **(Table 1)**. All extracts demonstrated antimicrobial activity against both fungi and bacteria. The largest inhibition zone against fungi was obtained when *P. aphidis* isolate L12 was extracted with heptane (2.0 cm^2^), followed by methanol and DCM (1.8 cm^2^), acetonitrile (1.6 cm^2^), EtOAc and MTBE (1.4 cm^2^). The smallest inhibition zone against fungi were obtained when *P. aphidis* was extracted with hexane and acetone (1.2 cm^2^). The largest inhibition zone against bacteria was obtained when *P. aphidis* was extracted with EtOAc (1.2 cm^2^), followed by heptane and hexane (1.1 cm^2^), methanol, acetone and acetonitrile (1.0 cm^2^) and DCM (0.9 cm^2^). The smallest inhibition zone against bacteria was obtained when *P. aphidis* was extracted with MTBE (0.7 cm^2^).

**Table 1.** Antimicrobial* activity of *P. aphidis* isolate L12 extractions with different organic solvents.

| **Extraction solvent** | **Antifungal activity**<br>**(Inhibition zone in cm<sup>2</sup>)** | **Antibacterial activity**<br>**(Inhibition zone in cm<sup>2</sup>)** |
| --- | --- | --- |
| **EtOAc** | **1.4** | **1.2** |
| **Methanol** | **1.8** | **1.0** |
| **Heptane** | **2.0** | **1.1** |
| **Hexane** | **1.2** | **1.1** |
| **MTBE** | **1.4** | **0.7** |
| **DCM** | **1.8** | **0.9** |
| **Acetone** | **1.2** | **1.0** |
| **Acetonitrile** | **1.6** | **1.0** |

\*Antifungal against *B. cinerea*. Antimicrobial against *A. tumefaciens*

Solvent-only controls did not cause any inhibition of neither fungi nor bacteria. Based on these results, we chose to continue our examinations and extract the metabolites from *P. aphidis* with EtOAc, due to the balanced between antifungal and antibacterial activity that we obtained with this solvent, its ability to dissolve compounds with wide range of polarity and the extensive use of this solvent in the extraction of natural compounds.

### Isolation of antimicrobial compounds from P. aphidis extract on TLC

*P. aphidis* isolate L12 extract was analyzed using bioautography with TLC plates, which enables natural products with antimicrobial activities to be localized in extracts (Hostettmann et al., 1997).

Followed the development of the plates, a total of 14 bends/metabolites were detected **(Figure 1)** and characterized according to their Rf and appearance under UV light (**Table 2**). Aoutobiography with *B. cinerea* indicated the presence of antimicrobial compounds in two areas (Figure 1; marked with arrows). As can be seen in **Figure 1** and **Table 2**, the location of the inhibition bands responds to the Rf of metabolites 11 (Rf 0.81) and 13 (Rf 0.93).

**Figure 1.**
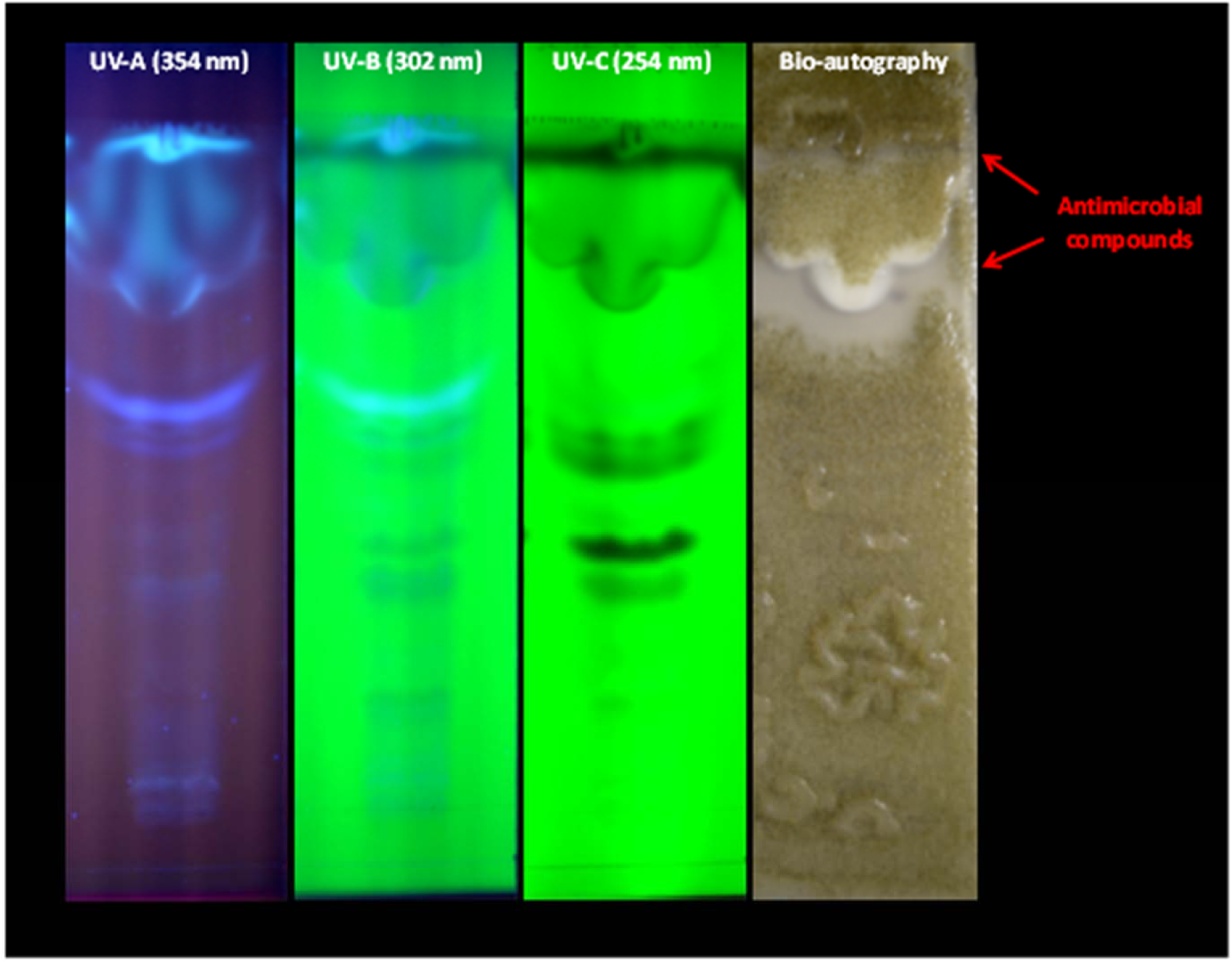
Thin-Layer Chromatography Bioautography of *P. aphidis* extract. After migration of *P. aphidis* extract on TLC plate, the plate was air dried and observed under UV light at three wavelengths: UV-A (354 nm), UV-B (302 nm) and UV-C(254 nm). Subsequently, the plate was covered with conidia suspension of B. cinerea (Bio-aoutography). The TLC plate was then incubated at 25°C, in darkness, for 7d. Metabolites with antifungal activity exhibit inhibition bands (marked with arrows).

**Table 2.**
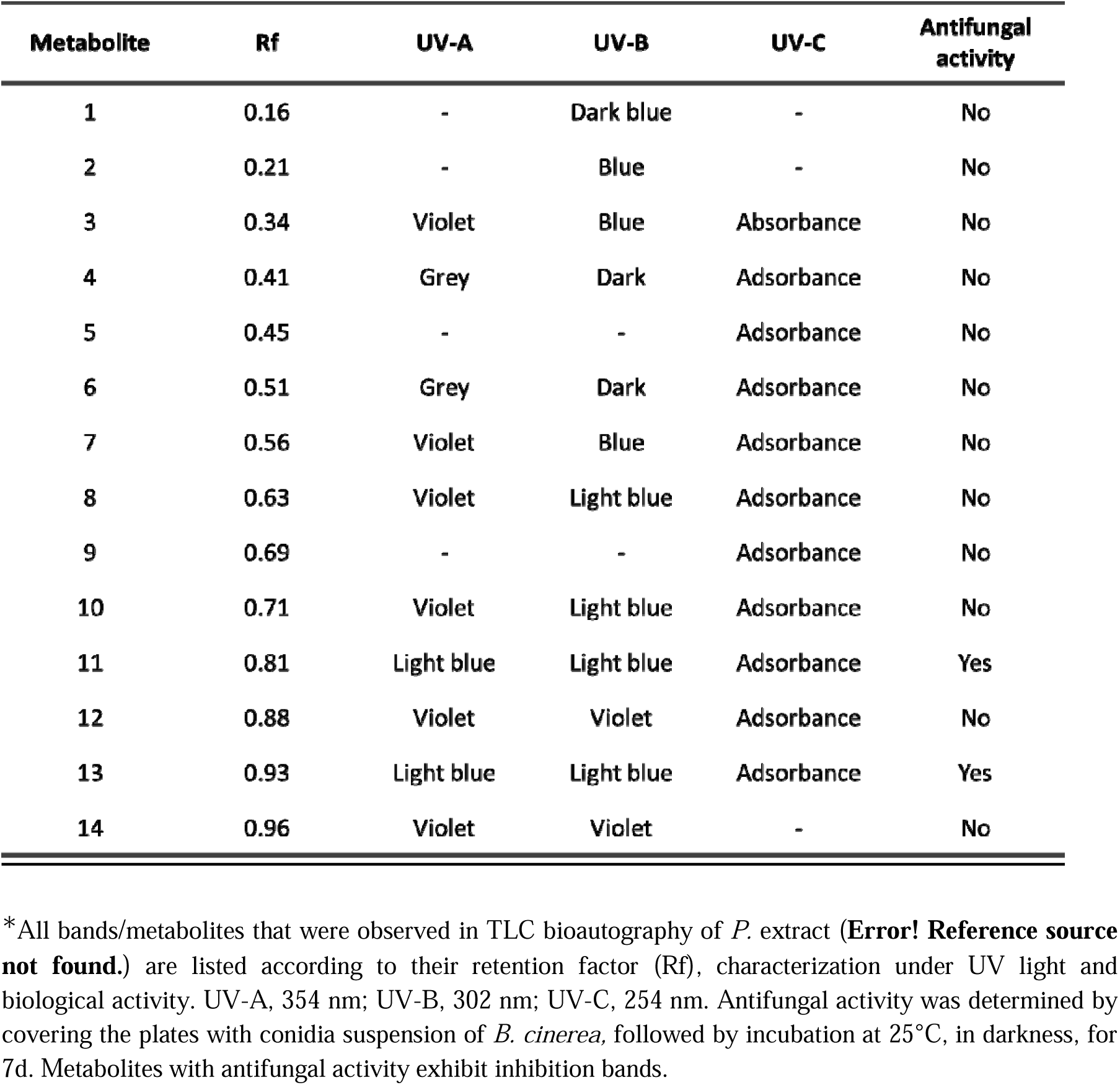
List of bands/metabolites of *P. aphidis* extract that were observed on TLC*.

### Isolation of the most active antimicrobial fraction using semi-preperative HPLC

The EtOAc extract was further fractionated with preparative RP-HPLC (**Figure S1**) into 11 fractions and all of the fractions were collected and subjected to agar diffusion assays to examine their antimicrobial activity against *B. cinerea* and *A. tumefaciens*. Only one fraction (number 2), was found active against both fungi and bacteria (**Table 3**). The obtained semi-purified active fraction was analyzed using high resolution HPLC-MS. Total ion chromatogram (TIC) of positive mode electrospray ionization (+ESI) scan showed 10 main chromatographic peaks (**Figure 2**). Further analysis of the fraction identified 11 compounds, all with relatively low molecular mass (<610 g mol^-1^). The molecular formula of two of the compounds (8 and 9 highlighted in blue) is characteristic to antibiotics (**Table 4**).

**Table 3.**
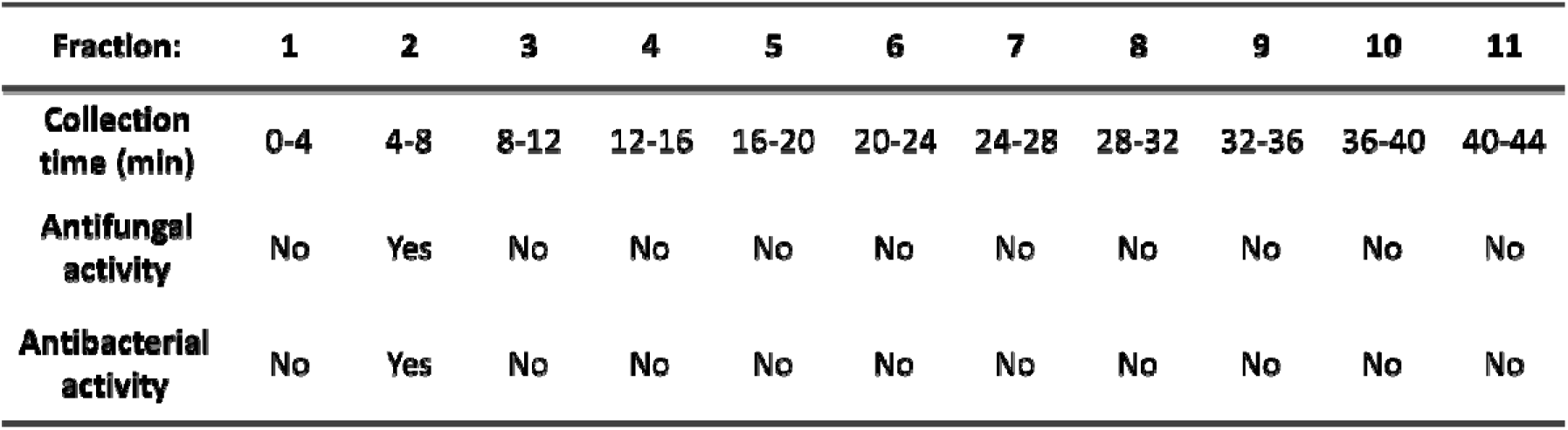
Antimicrobial activity of semi-purified fractions from *P. aphidis* isolate L12 extract.

**Figure 2.**
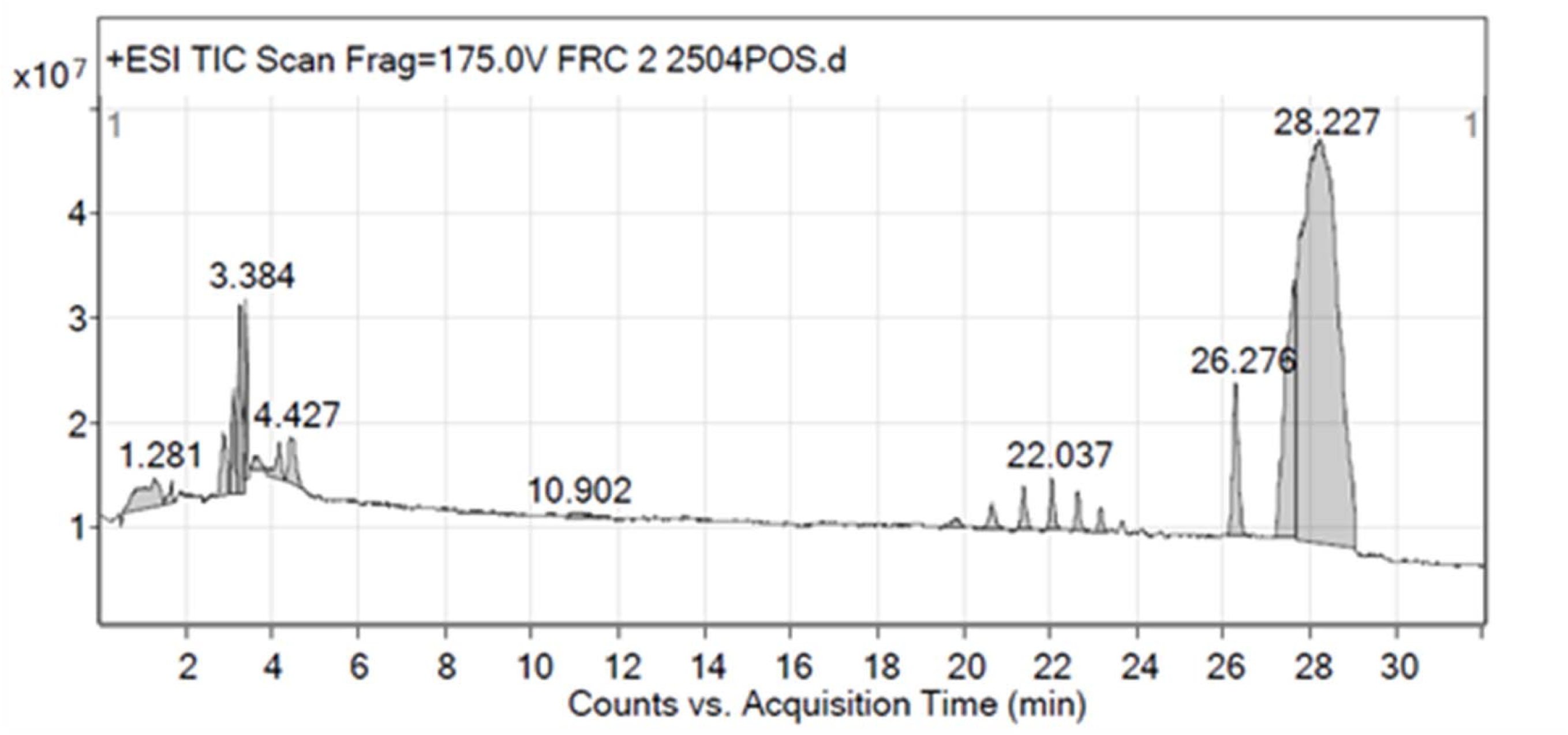
HPLC-MS analysis of *P. aphidis* antimicrobial fraction. Semi-purified and antimicrobial fraction of *P. aphidis* extract was analyzed with HPLC-MS . +ESI scan resulted in 10 chromatographic peaks.

**Table 4.**
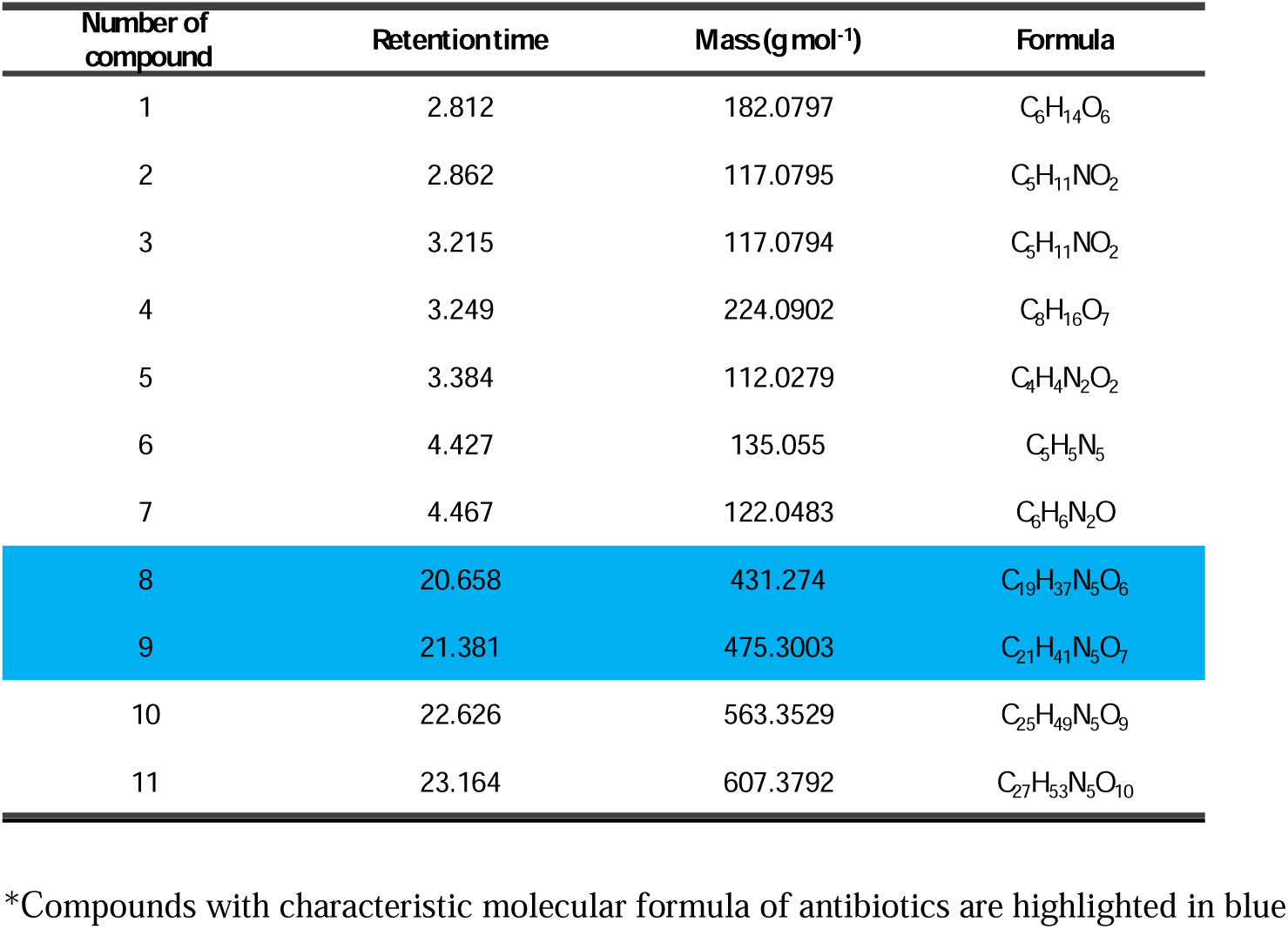
Identified compounds in *P. aphidis* isolate L12 semi-purified fraction, containing antimicrobial metabolites*.

### In vitro spore germination- and mycelial-inhibition of fungal phytopathogens

The inhibitory effect of *P. aphidis* extract on the spore germination of phytopathogenic fungi was demonstrated on several pathogens, such as *B. cinerea*, which causes gray mold and *A. alternata*, which causes leaf spot, rots and blights, and on a specialized pathogen - *Fusarium oxysporum f. sp. lycopersici* , which causes vascular wilt in tomato plants. All pathogens were inhibited by the EtOAc extract (**Figure 3**A). No inhibition zone was formed around disc saturated with Ethyl acetate as control. Additionally to the inhibitory effect of *P. aphidis* extract on the spore germination, it was further established that *P. aphidis* extract could also inhibit the mycelial growth of important plant pathogenic fungi. The mycelial growth-inhibitory effect was evaluated by calculating the IC50 values of the extract. The following IC50 values were obtained: 606µg ml^-1^ for *B. cinerea*, 221µg ml^-1^ for *Pythium spp*., 519µg ml^-1^ for *R. solani*, 455µg ml^-1^ for *S. sclerotiorum*, 2270µg ml^-1^ for *Fusarium oxysporum f. sp. Lycopersici*, and 2038µg ml^-1^ for *A. alternata* (**Figure 3B**).

**Figure 3.**
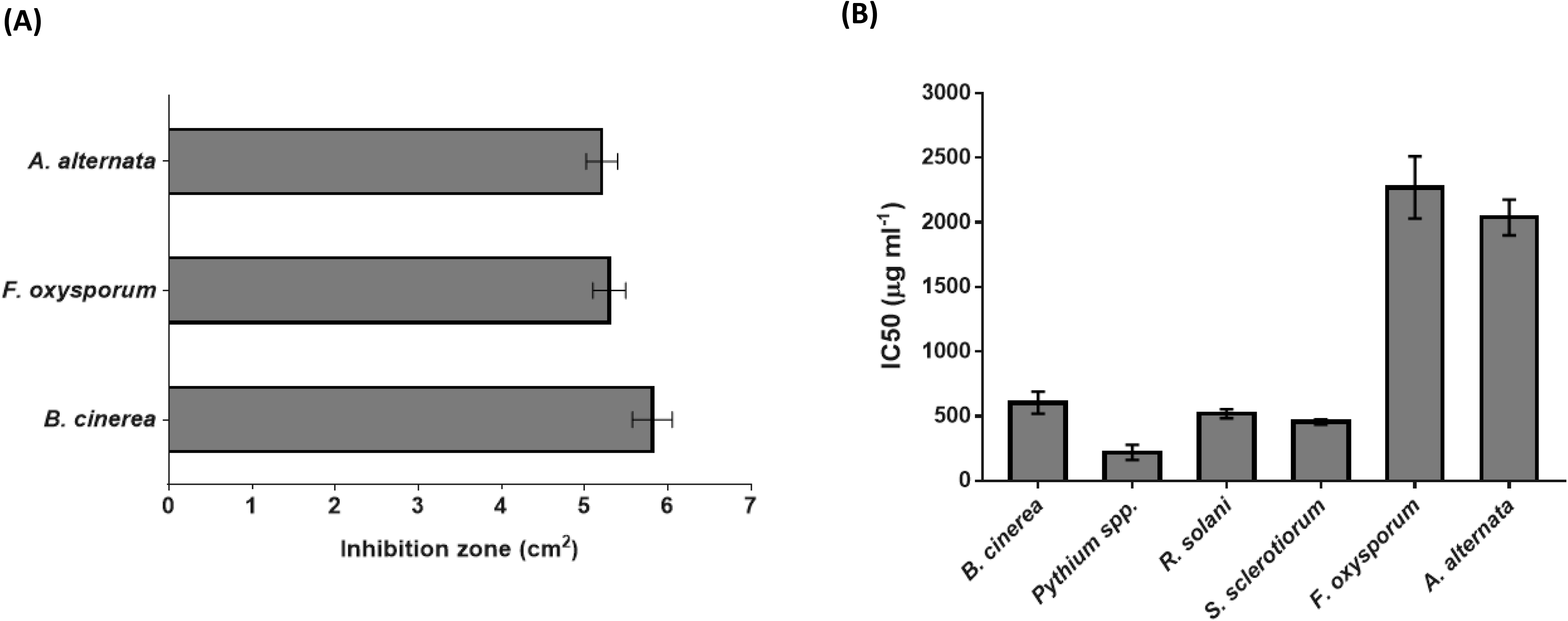
In vitro spore germination-and mycelial growth*-*inhibition of fungal phytopathogens by *P. aphidis* extract. **(A)** Spore inhibition. The area of the inhibition zone was measured 24 h after incubation of spores at the pathogen’s propagation temperature. *B. cinerea* (25°), *F. oxysporum* (27°) and *A. alternata* (25°). Results represent means ± standard error (n = 24). **(B)** Mycelial growth inhibition. Fungal mycelial plugs of *B. cinerea* (25°), Pythium spp.(27°) *R. solani* (27°), *S. sclerotiorum* (27°), *F. oxysporum* (27°) and *A. alternata* (25°) were placed on petri dishes, containing PDA culture media supplemented with increasing concentrations of *P. aphidis* extract. The IC50 values were calculated from the area of the treated colony, relative to the control (PDA only). Results represent means ± standard error (n = 12).

### In vitro growth inhibition of bacterial phytopathogens

The inhibitory effect of *P. aphidis* extract was also demonstrated on bacterial pathogens, such as *Pseudomonas syringae pv. tomato*, which causes bacterial speck, *X. campestris pv. Vesicatoria*, which causes bacterial leaf spot on pepper and tomato, *C. michiganensis subsp. michiganensis*, which causes bacterial wilt and canker of tomato, *E. amylovora*, which causes fire blight in apple and pear trees, and *A. tumefaciens*, which causes crown gall disease in a wide range of hosts plant. All bacterial pathogens that were examined were inhibited by the *P. aphidis* extract (**Figure 4**). No inhibition zone was formed around disc saturated with EtOAc as control.

**Figure 4.**
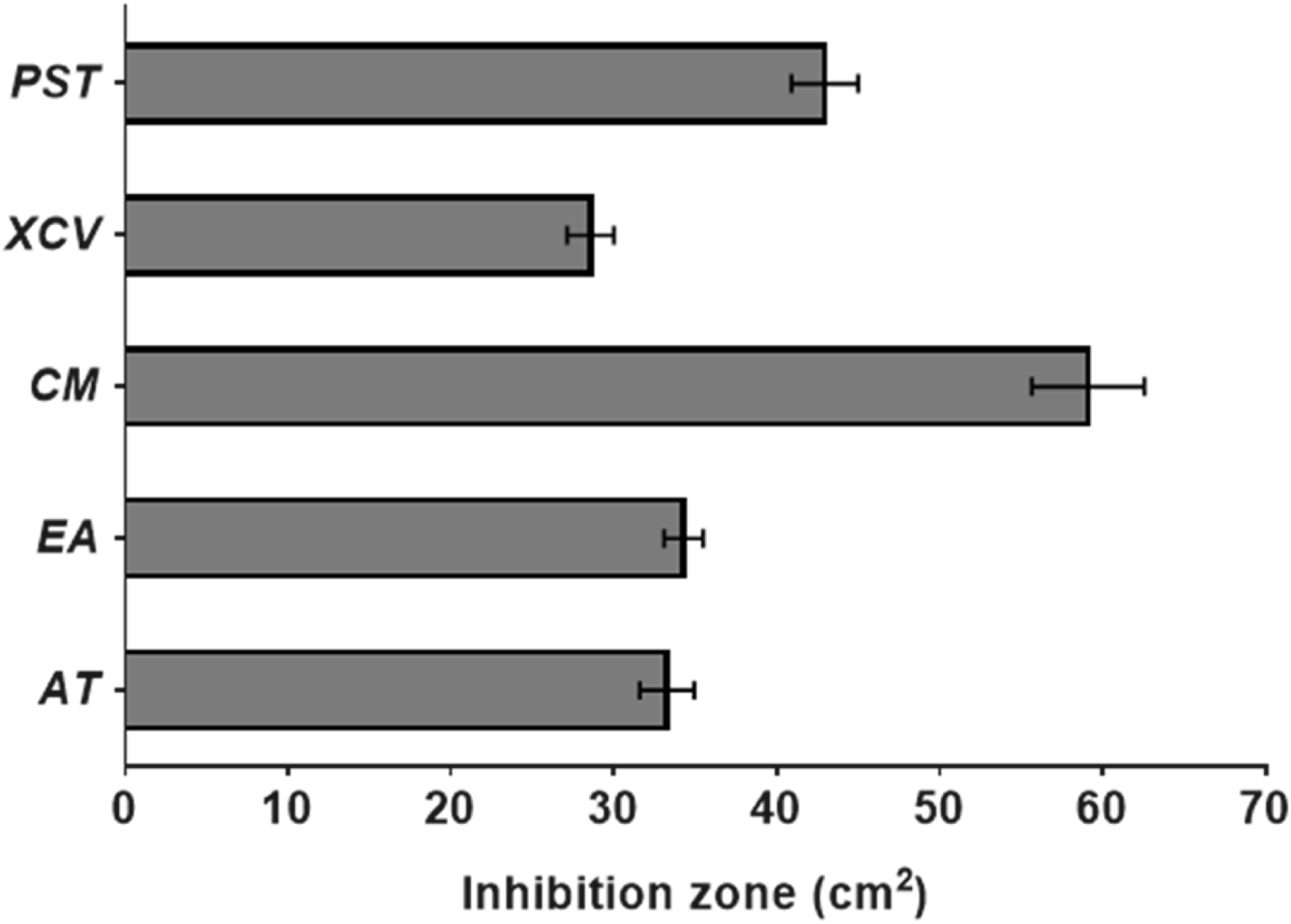
In vitro inhibition of bacterial phytopathogens by *P. aphidis* extracte. antimicrobial metabolites. The area of the inhibition zone was measured 24 h after incubation at the pathogen’s propagation temperature. PST = *P. syringe* pv. tomato, XCV = *X. campestris* pv. vesicatoria, CM = *C. michiganensis*, EA = *E. amylovora*, AT = A. tumefaciens. Results represent means ± standard error (n = 24).

### P. aphidis extract Inhibits B. cinerea Infection on tomato plants

Pre-treatment of the *B. cinerea* spores with the *P. aphidis* extract caused a dose-dependent reduction in the disease symptoms on detached tomato leaflets (**Figure 5A**). Pre-treatment with the extract in a concentration of 0.35 mg ml^-1^ reduced the area of the formed lesion to 78 mm^2^ (35% reduction), 0.7 mg ml^-1^ reduced the area of the formed lesion to 61 mm^2^ (49% reduction), 1.4 mg ml^-1^ reduced the area of the formed lesion to 42 mm^2^ (65% reduction), 4.2 mg ml^-1^ significantly reduced the area of the formed lesion to 15.5 mm^2^ (87% reduction) and 7 mg ml^-1^ significantly reduced the area of the formed lesion to 1.7 mm^2^ (98.5% reduction).

**Figure 5.**
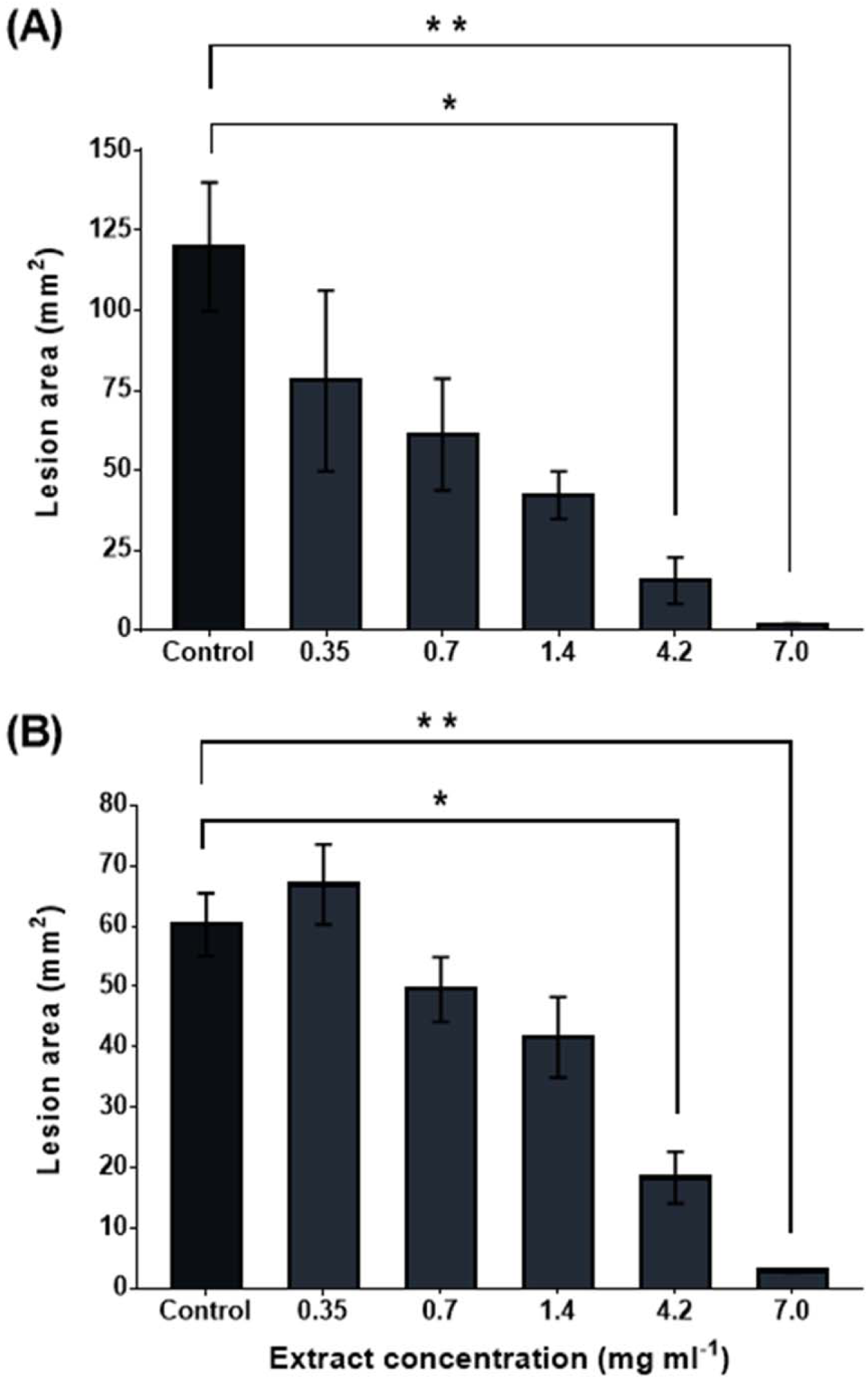
*P. aphidis* extract inhibit *B. cinerea* infection on tomato plants. Detached tomato leaflets (**A**), or intact tomato plants (**B**) were inoculated with *B. cinerea* spore suspension (1500 spores µl^-1^) pre-treated with increasing concentrations of *P. aphidis* extract, or 5% EtOAc (vol/vol) as control. The area of the *B. cinerea* lesions was monitored 96 h (A), or 120 h (B) after inoculation. Results represent means ± standard error (n = 6). Asterisks denote significant differences from control (P<0.05) as determined by one-way ANOVA and Dunnett’s multiple comparisons test. Shown here are the results of one experiment that was representative of at least three experiments with similar results.

Similar results were obtained with the intact tomato plant, although the progression of the disease symptoms was much slower than in the detached leaflets. 120 h post the inoculation, plants, which were inoculated with un-treated B. cinerea spores, developed necrotic lesions that covered a significant part of the leaflet (60 mm^2^). Pre-treatment of the *B. cinerea* spores with the *P. aphidis* extract caused a dose-dependent reduction in the disease symptoms (**Figure 5B**). Pre-treatment with the extract in a concentration of 0.35 mg ml^-1^ insignificantly increased the area of the formed lesion to 66 mm^2^ (10% increase), 0.7 mg ml^-1^ reduced the area of the formed lesion to 50 mm^2^ (17% reduction), 1.4 mg ml^-1^ reduced the area of the formed lesion to 42 mm^2^ (30% reduction), 4.2 mg ml^-1^ significantly reduced the area of the formed lesion to 18 mm^2^ (70% reduction) and 7 mg ml^-1^ significantly reduced the area of the formed lesion to 3 mm^2^ (95% reduction).

In both cases, detached leaflets and intact plants, significant reduction of the disease symptoms was obtained when the spore suspension was treated with extract concentrations higher than 4.2 mg ml^-1^ (**Figure 5A and B**). Higher concentration than 14 mg ml^-1^ of this extract caused significant visible damage to the plants (data not shown). The preventive effect of *P. aphidis* extracted antimicrobial metabolites was evaluated by spraying the plants with increasing concentrations of the extract several hours before the inoculation of the plants with *B. cinerea*. 7 days post inoculation, the untreated plants developed severe symptoms on most of the inoculated leaflets (81% infected leaflets). Pre-treatment of the plants (spraying) with *P. aphidis* extract in concentrations of 0.2 mg ml^-1^ (73% infected leaflets) and 1 mg ml^-1^ (84% infected leaflets) did not have significant inhibitory effect on the levels of the *B. cinerea* infection. However, pre-treatment of the plants with *P. aphidis* extract in concentrations of 5 mg ml^-1^ (35% infected leaflets), 10 mg ml^-1^ (34% infected leaflets) and 15 mg ml^-1^ (26% infected leaflets) led to a significant reduction in the portion of the inoculated leaflets, which developed disease symptoms (**Figure 6**).

**Figure 6.**
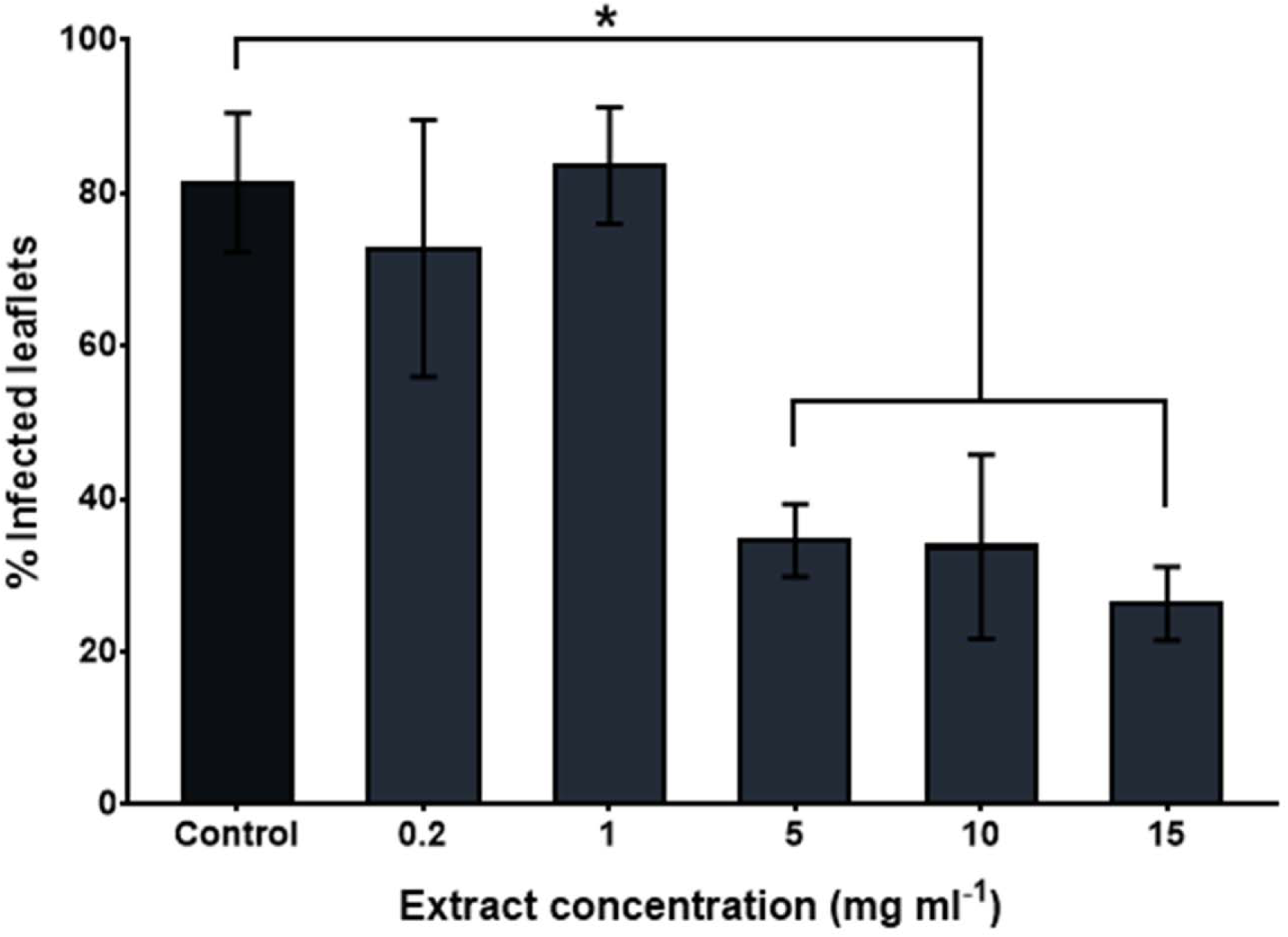
Preventive effect of *P. aphidis* extract against *B. cinerea* infection. Tomato plants were sprayed with increasing concentrations of *P. aphidis* extract leaflets were then inoculated with B. cinerea spore suspension. The disease symptoms were evaluated after 7 days as the percentage of infected leaflets per plant. Results represent means ± standard error (n = 5). Asterisks denote significant differences from control (P<0.05) as determined by one-way ANOVA and Dunnett’s multiple comparisons test. Shown here are the results of one experiment that was representative of at least three experiments with similar results.

Characterization of induced resistance obtained by *P. aphidis* aqueous extract *in planta*.

Bioassay-guided isolation of a semi-purified aqueous fraction of *P. aphids*, did not demonstrate direct antimicrobial activity (**Figures S2 and S3**), this allowed us to distinguish the effects of IR activation from the effects of direct antimicrobial activity.

### Local induction of defense-related genes by the semi-purified aqueous extract of Pseudozyma aphidis

To verify the ability of our semi-purified aqueous fraction to activate plant resistance, the local effect of application of the semi-purified aqueous fraction on the expression of defense-related genes wrer evaluated on tomato plants.

The expression of the four defense related genes was up-regulated during the first 24 h after application of the semi-purified aqueous fraction on the plants, peaked around 24 h after application and was moderately down-regulated over the next 72 h (**Figure 7**). 120 h post application, the relative expression of the genes was still significantly higher than the basal expression.

**Figure 7.**
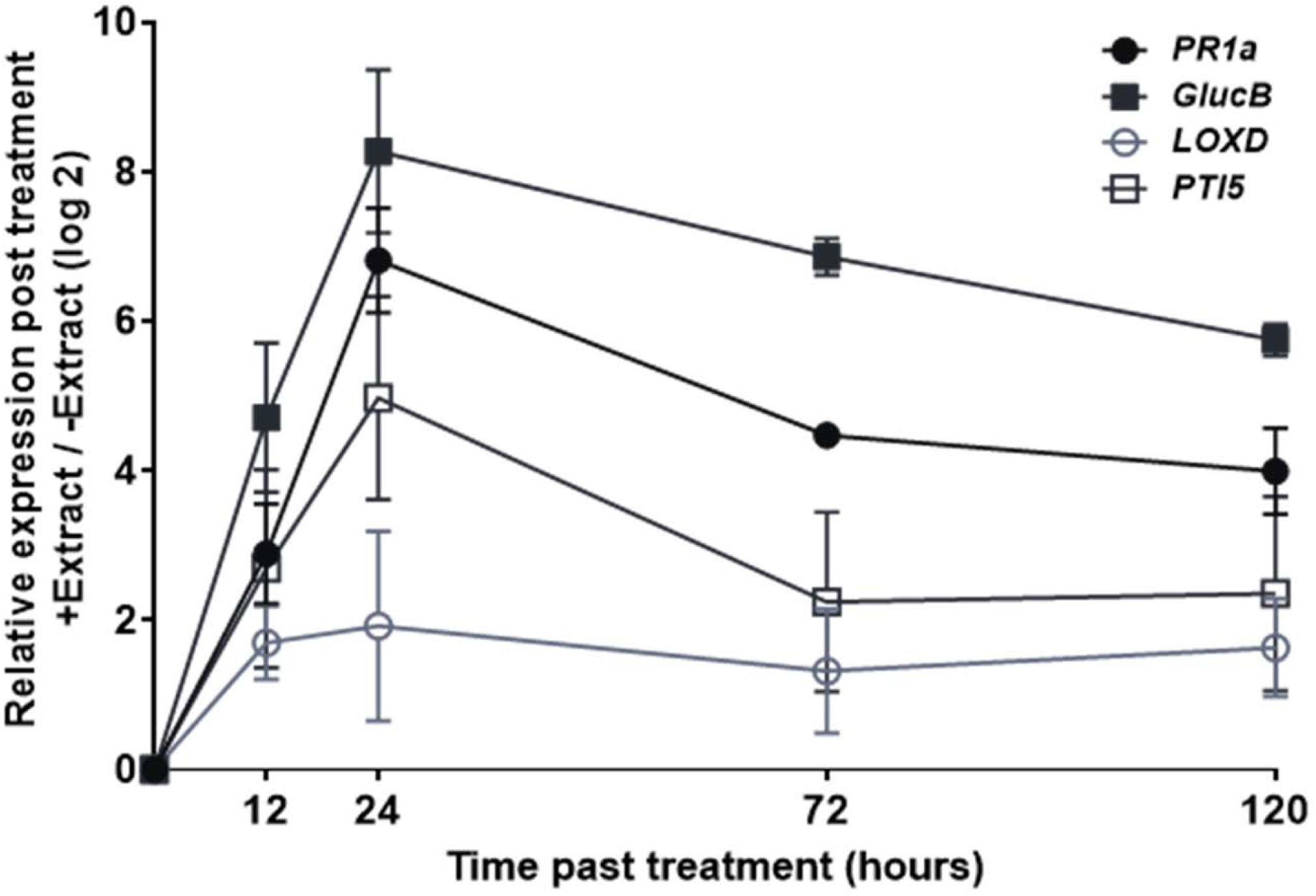
Local induction of defense-related genes by semi-purified aqueous fraction of *P. aphidis* extract. Tomato plants were sprayed with the semi-purified aqueous fraction and gene expression was ,easured 12, 24, 72 and 120 h after treatment. Relative gene expression was calculated by the comparative cycle threshold 2^-ΔΔCt^ method. Results represent means ± standard error (n=3). Shown here are the results of one experiment that was representative of at least three experiments with similar results.

In spite of the similar trend in the effect of the semi-purified aqueous fraction on the different genes, there was a difference in the intensity of their expression. The expression of *PR1a* and *GlucB*, which are related to the SA pathway were up-regulated more than 100 fold, while the expression of *LOXD*, which is related to the JA pathway was up-regulated less than 10 fold.

### Systemic induction of defense-related genes by semi-purified aqueous extract of Pseudozyma aphidis

The effect of application of the semi-purified aqueous fraction on the expression of the defense-related genes was examined also systemically (**Figure 8A**). The results did not show a significant difference between the “short distance” systemic effect (proximate leaflets; **Figure 8A**) and the “long distance” systemic effect (systemic leaflets on other leaves; **Figure 8B**). On both cases, the expression of *GlucB* and *LOXD* was rapidly up-regulated during the first 24 h, and then stayed relatively constant for the next 72 h. In contrast, *PR1a* and *PTI5* were considerably less systemically affected by the application of the semi-purified aqueous fraction. The expression of *PTI5* was weakly up-regulated during the first 12 h and then was quickly down-regulated back to the basal level. The expression of *PR1a* was weakly down-regulated during the first 72 h before coming back to the basal levels. In comparison to the results of the local induction, it is interesting to notice that while the application of the semi-purified aqueous fraction locally induced the expression of the all four defense-related genes, only two of the genes were systemically affected in a similar way.

**Figure 8.**
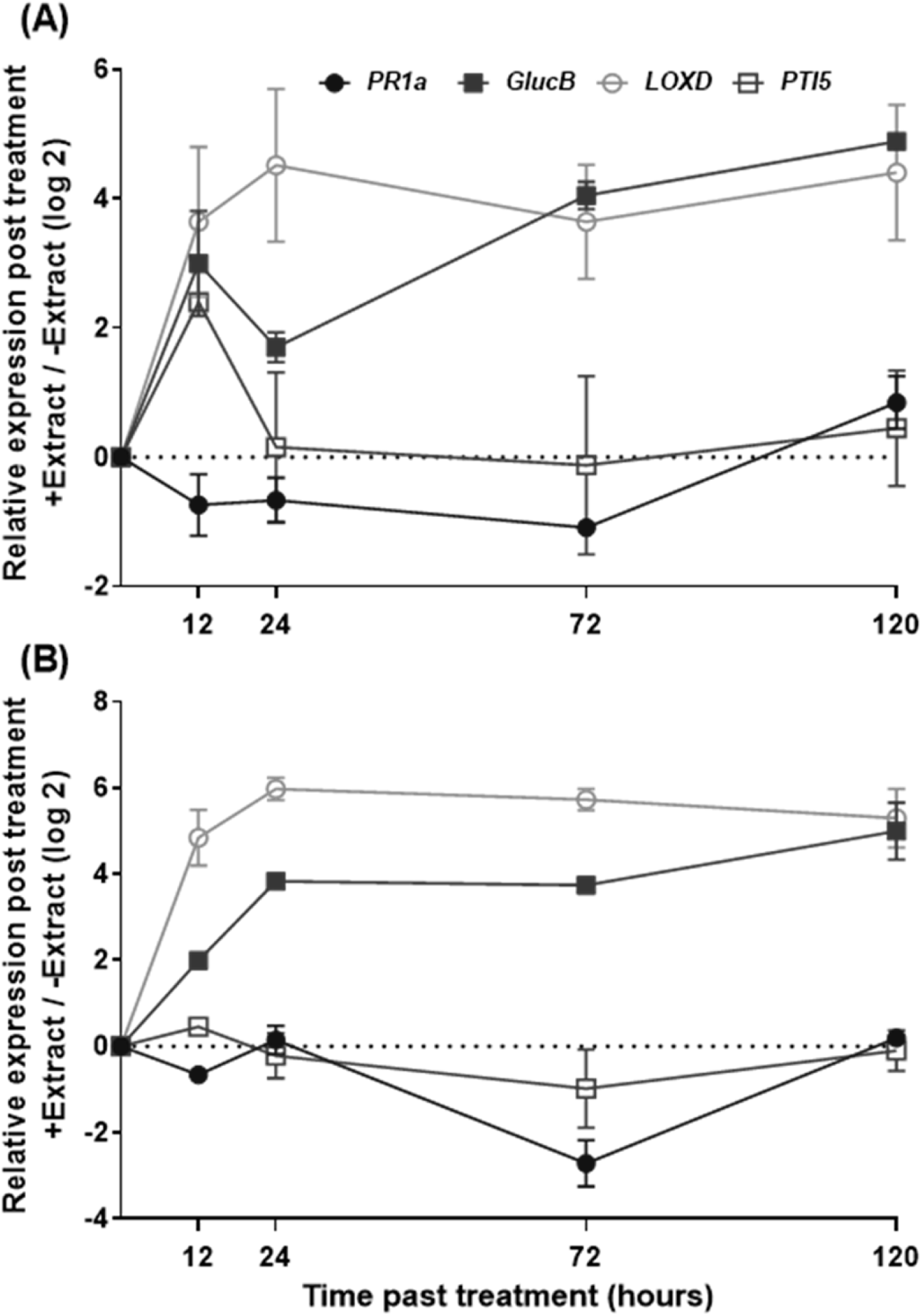
Systemic induction of defense-related genes by semi-purified aqueous extract of *P. aphidis*. Samples were taken 12, 24, 72 and 120 h after the treatment from proximate leaflets on the same leaf (**A**), or systemic leaflets on another leaf (**B**). Relative gene expression was calculated by the comparative cycle threshold 2^-ΔΔCt^ method. Results represent means ± standard error (n=3). Shown here are the results of one experiment that was representative of at least three experiments with similar results.

## Discussion

For several decades, constant use of chemical-based pesticides allowed farmers to control the spread of pathogens and pests and helped to ensure stable and prosperous agricultural systems. In recent years, this had started to change with a continuous rigorousness of the regulations against synthetic pesticides, due to human health and environmental concerns. However, for this trend to expand globally and bring a significant reduction in the total use of synthetic pesticides, reliable and cost-effective alternatives are required. One of the most promising alternatives is the discovery and developing of new BCAs and natural-based pesticides (Copping & Duke, 2007, Dayan et al., 1999, Dayan et al., 2009, Denholm & Rowland, 1992b, Leroux et al., 2002, Zhang et al., 2023, Daraban et al., 2023, Singh & Maurya, 2023).

Isolation of natural compounds from microorganisms is a challenging task, which requires a combination of different chromatographic, analytic and microbiological techniques. The classical way of isolation of natural compounds starts with identification, collection and preparation of the biological material. Extraction with different solvents from low to higher polarity follows. Isolation of semi-purified fractions, containing the biologically active compounds is achieved using bioassay-guided isolation strategies that connect information on the chemical profiles of extracts and fractions with their activity data in in vitro bioassays (Li *et al*., 2024, Majik & Gawas, 2023, Simoben *et al*., 2023).

Previous works in our laboratory demonstrated that *P. aphidis* (isolate L12) can reduce *B. cinerea* infection of tomato and Arabidopsis plants, by both direct antagonistic interactions and induced resistance (Buxdorf et al., 2013a, Buxdorf et al., 2013b). Later work in our laboratory also demonstrated that *P. aphidis* has an inhibitory effect on *C. michiganensis* in planta (Barda et al., 2015). Here we examined and characterized the bio-active compounds, produced and stored by this isolate.

*P. aphidis* biomass was extracted with several organic solvents with different hydrophobicity. Relatively similar antifungal and antimicrobial activity was measured in all the obtained extracts (**Table 1**), which indicates that the antimicrobial metabolites have intermediate polarity. Polar and highly hydrophilic compounds, which are dissolved in the cell’s cytoplasm would not have been extracted into the non-polar solvents such as heptane and hexane(Ito, 2005, Snyder, 1978). It is also possible that *P. aphidis* produces several antimicrobial compounds with different degrees of hydrophobicity. EtOAc was chosen for further preparation of *P. aphidis* extract. This organic solvent has intermediate hydrophobicity together with relatively high polarity, which enable it to dissolve and extract a very wide range of compounds. Due to those properties EtOAc is commonly used as an extraction solvent of natural compounds (Tamokou *et al*., 2012, Ogunnupebi *et al*., 2020, Singh *et al*., 2023).

TLC analysis of the *P. aphidis* extract confirmed that it contains at least 14 different metabolites, from which at least two have antifungal activity (**Figure 1**; **Table 2**). The absolute separation of the bands means that at least two different antimicrobial compounds originated them. It is important to remember that while each of the bands indicates the presence of an antimicrobial compound, the relatively low resolution of this technique doesn’t enable to determine whether each inhibition band is originated from only one antimicrobial compound, or several antimicrobial compounds with a similar chemical structure (Poole, 2003). The proximity of the two bands also suggest that even if there are more than two antimicrobial compounds, all of them have relatively similar chemical structure. Many techniques had been published for the isolation of bio-active fractions, with the main separation technologies used in recent years are methods of liquid chromatography such as HPLC. Natural products are often present only as minor component in the extract and the resolving power of HPLC is ideally suited to the rapid processing of such multicomponent samples on both an analytical and preparative scale (Sasidharan *et al*., 2011). HPLC fractionation of *P. aphidis* EtOAc extract revealed only one fraction that contain antimicrobial activity (**Figure S1 and Table 3**). Like the results of the TLC analysis, this suggests that all the antimicrobial compounds have similar chemical structure. Further analysis of the active fraction with high-resolution MS detector revealed that the semi-purified fraction contained 11 compounds, from which two compounds have molecular mass that is characteristic to antibiotics (Figure 2 **and Table 4**).

The inhibitory effect of the EtOAc extract was demonstrated against a range of fungal and bacterial pathogens (**Figures 3-6**). The results of the *in vitro* experiments demonstrate that the extracted metabolites are well capable of inhibit all the fungi that were examined. However, while the inhibitory effect on the spore germination was similar for all the examined fungi (**Figure 3A**), the inhibitory effect on the mycelial growth was much more selective (**Figure 3B**). While the inhibitory effect on the spore germination was almost equal for *B. cinerea, F. oxysporum* and *A.alternata*, the inhibitory effect on the mycelial growth of *F. oxysporum* and *A.alternata* was much weaker than the inhibitory effect on the mycelial growth of other fungi (**Figure 3B**).

*In vitro* results of bacteria inhibition by the extracted metabolites demonstrate strong activity against all tested bacteria (**Figure 4**). Antimicrobial activity of fungal secondary metabolites is well documented in the literature (Mousa & Raizada, 2013).

Our results are also correspondent with the results of Barda et al. (2015)(Barda et al., 2015), who performed a similar experiment with an extraction of the *P. aphidis* filtrate. Interestingly, although lower amounts of the extract were used in the present experiment, the inhibition of similar bacteria was much stronger than in the experiment performed by Barda et al., 2015(Barda et al., 2015). This indicates that the concentration of the active metabolites in the biomass extract is much higher than in the filtrate, and may suggest that most of the active metabolites which are produced by *P. aphidis*, are stored in the cells, and secreted to the medium in relatively small amounts.

Fungi inhibition by *P. aphidis* extract was also demonstrated in planta, against the important airborne plant pathogen *B. cinerea*, which causes gray mold in almost all major greenhouse crops (Rosslenbroich & Stuebler, 2000, Leroux, 2007, Elad *et al*., 2007, Fillinger, 2016, Williamson *et al*., 2007). Gray mold is one of the most common diseases of greenhouse tomato, causing among the rest, visible symptoms on infected leaves (Williamson et al., 2007, van Kan, 2006, O’Neill *et al*., 1997). Results showed that application of *P. aphidis* EtOAc extract could reduce gray mold infection in tomato plants. Similar results were obtained for detached tomato leaflets and intact tomato plants (**Figure 5A and B**). Both experiments demonstrated a dose-dependent reduction in the infection. A reduction of over 95% in the infection symptoms was observed when the *B. cinerea* spores were treated with an extract concentration of 7 mg ml-1, and a significant preventive effect was observed when the *P. aphidis* extract was applied in concentration higher than 5 mg ml-1 (**Figure 6**). Only concentrations twice as high (14 mg ml-1) caused significant visible damage to the plants, probably by inducing programmed cell death (data not shown). Our recent results demonstrate the ability of *P. aphids* secretions to induce reactive oxygen species production and programmed cell death in *B. cinerea* (E. calderon et al., 2018), this can also indicate the ability of our extract to induces programmed cell death in plant tissue.

The inhibitory effect on *B. cinerea* disease on tomato plants also agree with similar studies that were previously conducted in our laboratory with the live spores of *P. aphidis*. Application of *P. aphidis* live spores on detached tomato leaflets and intact tomato plants three days before inoculation with *B. cinerea*, reduced the number of infected leaflets by more than 80% in the detached leaflets and up to 75% in the intact plants (Buxdorf et al., 2013). The strong correlation between the in planta inhibition activity of live *P. aphidis* and the in planta inhibition activity of *P. aphidis* extract, combined with our in vitro results, strongly supports our hypothesis that antibiosis is one of the main modes of action of *P. aphidis* (Buxdorf et al., 2013b, Buxdorf et al., 2013a, Barda et al., 2015, Calderón *et al*., 2018).

The ability of BCAs to activate IR in plants is well established in the literature, either before pathogen attack (Shoresh *et al*., 2010a, Nguvo & Gao, 2019, Srivastava *et al*., 2021), or by priming the induced defense (Conrath *et al*., 2002, Conrath *et al*., 2015, Molinari & Leonetti, 2019, Reddy, 2013). For example, *Trichoderma harzianum* T39 primed grapevines against downy mildew (Perazzolli *et al*., 2011) and *T. asperellum* SKT-1 induced resistance in Arabidopsis against the bacterial pathogen *Pseudomonas syringae* pv. tomato (Yoshioka *et al*., 2012, Schumacher, 2012) and modulated the expression of genes involved in the JA/ET-signaling pathways of ISR (Shoresh *et al*., 2005, Shoresh *et al*., 2010b). The endophytic basidiomycete fungus *Piriformospora indica* primed various plant species against powdery mildew (Molitor *et al*., 2011) and the protective *F. oxysporum* strain Fo47 primed tomato plants against pathogenic *F. oxysporum* strains (Aimé *et al*., 2013). The potential of mimicking this effect using extracted metabolites was also demonstrated (Djonović *et al*., 2006, Mehari *et al*., 2015). The signal transduction mechanisms that are involved however still need to be further investigated.

To thoroughly investigate the signal transduction pathways that are activated by *P. aphidis* extracted metabolites, the expression of defense-related genes was measured after application of the semi-purified aqueous fraction of *P. aphidis* extract. Our results demonstrate that application of the semi-purified aqueous fraction on tomato plants rapidly up-regulated the expression of most of the defense-related genes in the plant’s tissue that came with direct contact with the fraction (**Figure 7**). The induced expression had a parabolic trend, which peaked 12-48 h after the application. This result was repeated and verified in a more extensive experiment with four of the genes (*PR1a, GlucB, LOXD and PTI5)*, which represent both the ISR and the SAR pathways (**Figure 7**). The similar induction of genes that represent both pathways may indicate that *P. aphidis* metabolites can elicit IR via a complex pathway, which doesn’t fall under the clear definition of either the ISR or the SAR. These results do not fit with the traditional definition of IR in the literature, which divided into two different pathways, ISR and SAR, which work separately and even antagonistically toward each other (Kamle *et al*., 2020, Anderson *et al*., 2004). However, more recent works have established that in practice, many times this is not the case and the signal transduction pathways which are involved are more complex (Thomma *et al*., 2001, Robert-Seilaniantz *et al*., 2011, Li *et al*., 2019, Pieterse & Van Loon, 2004, Pieterse *et al*., 2012).

When the expression of the same representative genes was examined systemically (either in proximate leaflets or in systemic leaflets on other leaves), two of the genes (*GlucB and LOXD)* were still significantly affected and followed a similar trend (**Figure 8A and B**). This is especially important since a major advantage of the ISR and SAR is their systemic effect. While the local activation of the plant’s induced resistance can serve as a complementary mode of action to the direct antimicrobial metabolites of *P. aphidis*, activation of the plant’s systemic defense adds an additional layer of defense.

Our results are similar to the results of Djonović et al. (2006)(Djonović et al., 2006), who demonstrated that application of a secreted elicitor from *T. virens* locally and systemically up-regulated defense-related genes (*PR*, *Gluc*, *CHI* and *LOX1)* in cotton, associated with both the ISR and SAR pathways. Mehari et al. (2015)(Mehari et al., 2015) also demonstrated the up-regulation of similar tomato defense-related genes (*PR1a*, *Chi9* and *GlucB*), using biochar as an elicitor for IR.

Although the induction of marker genes is indicative of the induced pathway, additional experiments and results are still necessary for an accurate mapping of the pathways that are involved in the elicitation of IR and priming by *P. aphidis* extracted metabolites. These include a complete sequencing of the plant’s transcriptome after application of the semi-purified aqueous fraction and the use of plant mutants in hormonal pathways. None the less, our findings that *P. aphidis* may elicit IR via a non-traditional pathway is strongly supported by previous studies of our laboratory, which involved the use of tomato and *Arabidopsis* mutant plants with impaired SA accumulation and *Arabidopsis* mutants impaired in JA signaling. A comparison of our results to those studies also shows good correlation between the IR activation by *P. aphidis* extracted metabolites and the IR activation by *P. aphidis* isolate L12 live spores. In great similarity to our results, Barda et al. (2015)(Barda et al., 2015) found that application of *P. aphidis* live spores on tomato plants induced the expression of several defense-related genes of the SA and the ET pathways, including *PR1a, GlucB* and *PTI5*. These results also correspond with the induction of *PR1* in *Arabidopsis* plants following the application of *P. aphidis* live spores, which observed by Buxdorf et al. (2013). Similarly, to our results, Buxdorf et al. (2013) also observed activation of the JA pathway by the induction of *PDF1.2* gene after application of *P. aphidis* live cells(Buxdorf et al., 2013a). Contradictory, Barda et al. (2015) found no induction of LOXD, which represent the JA pathway after application of *P. aphidis* live cells. Both Buxdorf et al. (2013) and Barda et al. (2015) also observed priming of the plants’ IR after application of *P. aphidis* live cells(Barda et al., 2015, Buxdorf et al., 2013a). To our knowledge, only one other research group recently published the elicitation of IR and priming by *Pseudozyma* spp. live cells (Lee *et al*., 2017)and there are no publications regarding the extraction of IR elicitors from *Pseudozyma* spp.

The good correlation between the IR activation by *P. aphidis* extracted metabolites and the IR activation by *P. aphidis* live spores suggests that *P. aphidis* elicits this activity using low molecular weight metabolites, which can be extracted and isolated. The lack of direct antibiosis activity by the semi-purified aqueous fraction means that different sets of metabolites are responsible for the antibiosis and the IR. Therefore, the extracted IR elicitors can be used separately, or together with the antimicrobial compounds as part of natural-based pesticides with a dual mode of action.

## Author Contributions

Conceptualization, M.L. and R.H.; methodology, M.L. and R.H..; software, M.L. and R.H.; validation, M.L. and R.H. investigation, R.H.; writing—original draft preparation, M.L. and R.H.; visualization, M.L. and R.H.; supervision, M.L.; project administration, M.L.; funding acquisition, M.L. All authors have read and agreed to the published version of the manuscript.

## Data Availability Statement

Data supporting reported results can be found on HUJI data base cloud and upon re-quest.

---

**Figure S1.** Preparative RP-HPLC fractionation of P. aphidis extract. P. aphidis extract was fractionated with preparative RP-HPLC. A total of 11 fractions were collected.

**Figure S2.** Absence of antimicrobial activity by the semi-purified aqueous fraction of *P. aphidis* extract. Samples from EtOAc (**A**) or aqueous extract (**B**) were used in diffusion assays against the fungal phytopathogen *B. cinerea*. The area of the inhibition zone was examined 24 h after incubation at 22°C.

**Figure S3.** Absence of antimicrobial metabolites in the semi-purified aqueous fraction of *P. aphidis* extract. After migration, the TLC plates were air dried and observed under UV light at two wavelengths: UV-A (354 nm) and UV-C(254 nm). Migrated metabolites that previously exhibited inhibition bands (see Figure 1) are visibly absence in the semi-purified aqueous fraction (AF), compared to the organic fraction (OF) and the unfractionated *P. aphidis* extract (Ext).

