## supplementary figures and methods for "Pseudozyma aphidis bio-active extract inhibit plant pathogens and activate induce resistance in tomato plants"

###### SUPPLAMENTARY Haris and Levy

###### Table S1. Primers used in the qRT-PCR analysis

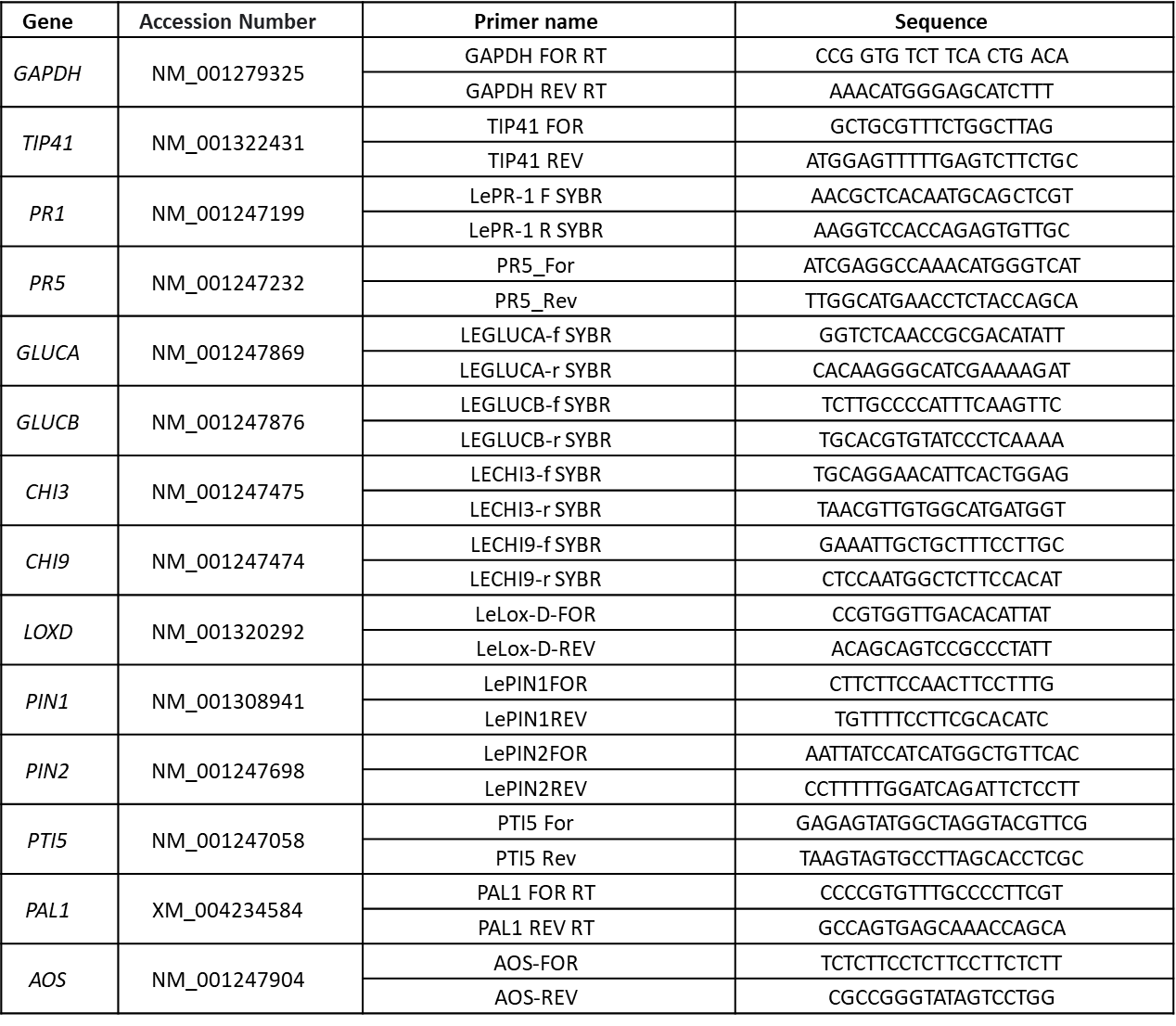

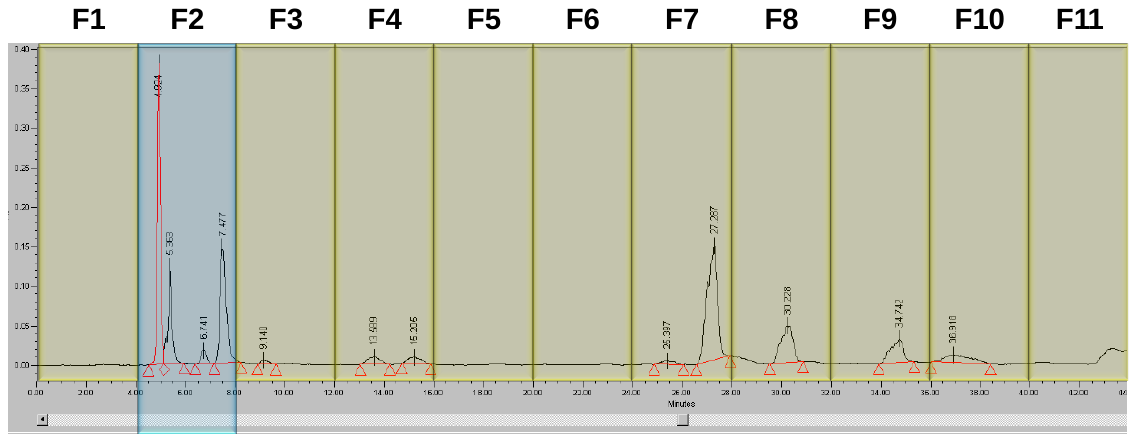

#### Figure S1. Preparative RP-HPLC fractionation of P. aphidis isolate L12 extract.

### *P. aphidis* isolate L12 extract was fractionated with preparative RP-HPLC. A total of 11 fractions were collected.

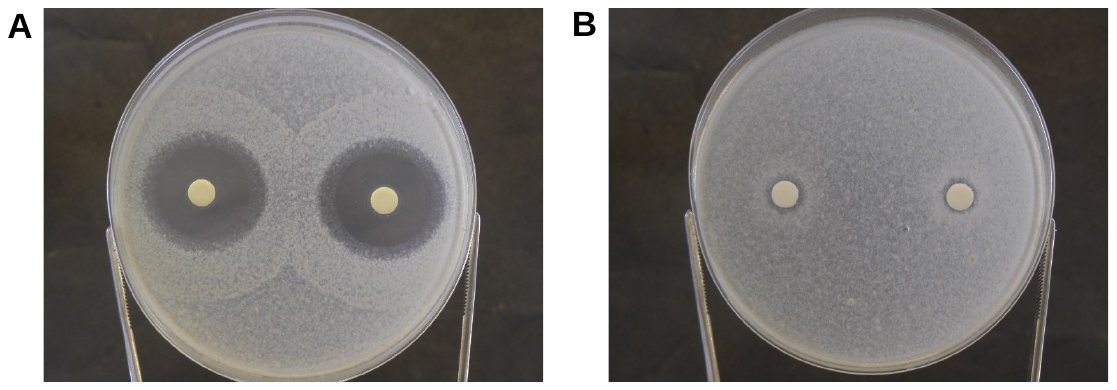

#### Figure S2. Absence of antimicrobial activity by the semi-purified aqueous fraction of P. aphidis extract. 20 µl of a diluted sample, corresponding to 1 mg in dry weight, were used in diffusion assays against the fungal phytopathogen B. cinerea. The area of the inhibition zone was examined 24 h after incubation at 22°C. strong spore germination-inhibition was obtained by P. aphidis extract (A), compared to complete absence of germination-inhibition (same as control) by the semi-purified aqueous fraction (B).

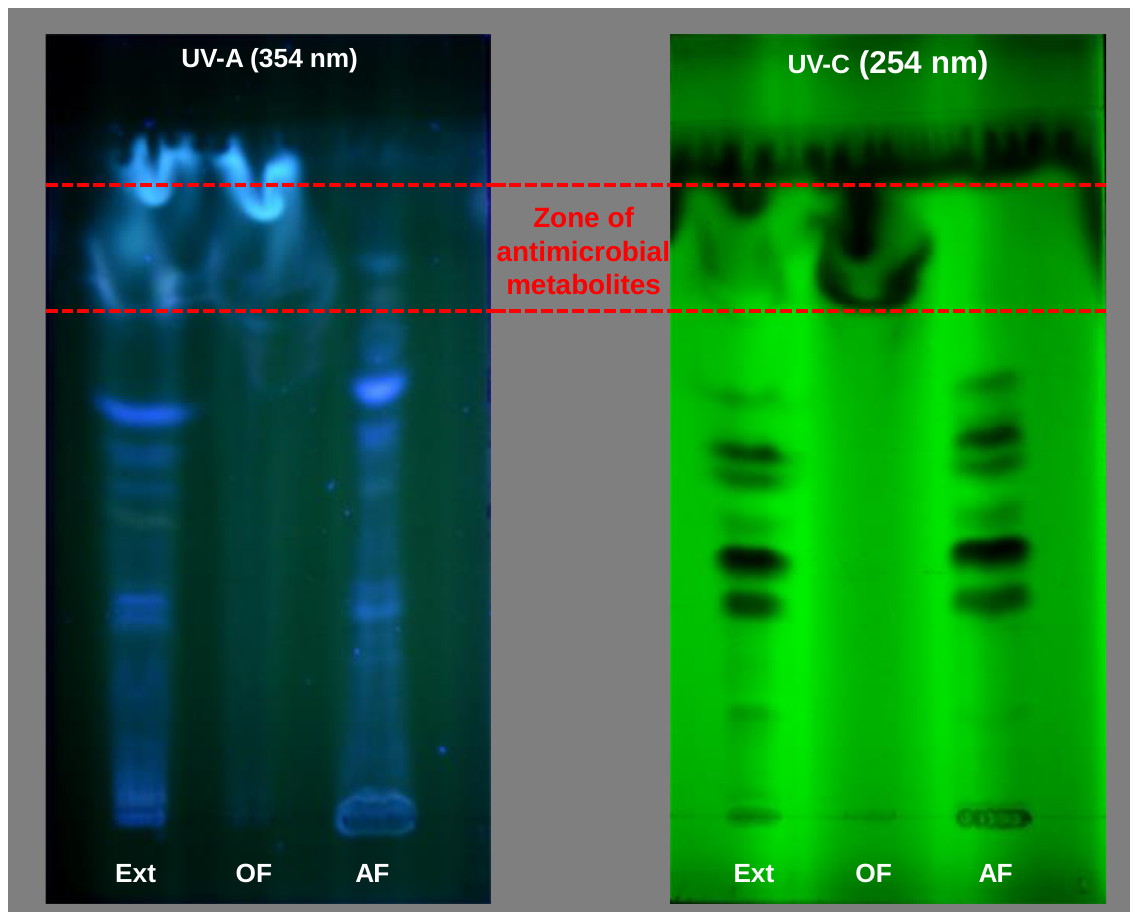

#### Figure S3. Absence of antimicrobial metabolites in the semi-purified aqueous fraction of P. aphidis extract.10 mg in dry weight, corresponding to 50 µl of sample were spotted on TLC plates (Silica gel 60 F_254_). The plates were developed under saturated conditions with the following solvent system: chloroform/methanol/water (75/25/2.5). After migration, the plates were air dried and observed under UV light at two wavelengths: UV-A (354 nm) and UV-C)254 nm(. Migrated metabolites that previously exhibited inhibition bands (see Figure 1) are visibly absence in the semi-purified aqueous fraction (AF), compared to the organic fraction (OF) and the unfractionated P. aphidis extract (Ext).
